# Efficient Pangenome Construction through Alignment-Free Residue Pangenome Analysis (ARPA)

**DOI:** 10.1101/2022.06.03.494761

**Authors:** Arnav Lal, Ahmed Moustafa, Paul J. Planet

**Author notes:** Corresponding author: Planet, Paul J.

## Abstract

Protein sequences can be transformed into vectors composed of counts for each amino acid (vector of Residue Counts; vRC) that are mathematically tractable and retain information about homology. We use vRCs to perform alignment-free, residue-based, pangenome analysis (ARPA; https://github.com/Arnavlal/ARPA). ARPA is 70-90 times faster at identifying homologous gene clusters compared to standard techniques, and offers rapid calculation, visualization, and novel phylogenetic approaches for pangenomes.

## Main Text

As millions of microbial genome sequences become available and whole genome epidemiology becomes the gold standard in infectious diseases, computationally tractable tools are needed for analysis of large genomic datasets. A critical step in comparative genomics is the construction of a pangenome, namely the union of genes within genomes (Medini *et al*. 2005). However, identifying gene “same-ness” across genomes, either as homology or orthology, is challenging—microbial pangenome construction is NP-Hard (Nguyen *et al*. 2014). Current conventional pangenomic methods rely on an alignment step, notably BLAST (Camacho *et al*. 2009), to compare genes, which requires additional steps such as pre-clustering, as performed in LS-BSR (Sahl *et al*. 2014) and Roary (Page *et al*. 2015), to simplify complexity. Alignment-free (AF) sequence comparison techniques using k-mer frequency, common substrings, micro-alignments or other heuristic or information theory techniques (as reviewed in Zeilezinski *et al*. 2017, 2019) are mostly focused on all-*vs*.-all pairwise whole genome distances with less emphasis on homologs or orthologs. Another AF technique, WhatsGNU (Moustafa *et al*. 2020) uses exact protein matches but cannot identify orthologs or homologs without accessing alignment-based techniques.

We developed an AF protein-based methodology, ARPA, to identify homologs, orthologs, and paralogs, and rapidly generate pangenomes. Instead of using amino acid alignments, we use twenty value vectors (vRC) composed of amino acid residue counts. To our knowledge, the ability to uniquely identify homologs using simple counts of amino acids is conceptually novel.

ARPA requires an input folder of “.faa” amino acid files. Encoded proteins are converted into a 1-by-20 row vector (vRC) with each term corresponding to the number of specific amino acid residues present within the sequence (Fig. 1A).

**Figure 1.**
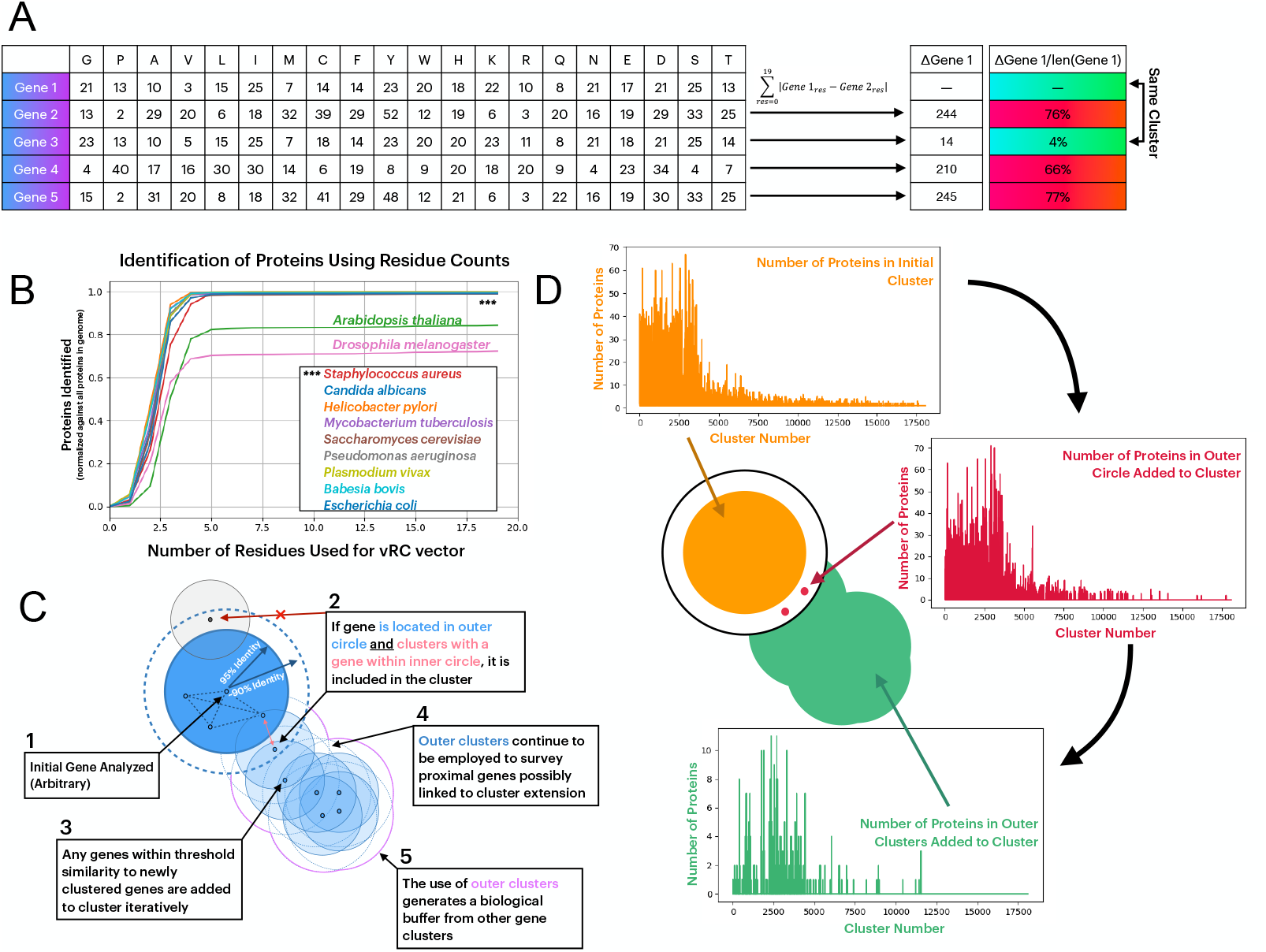
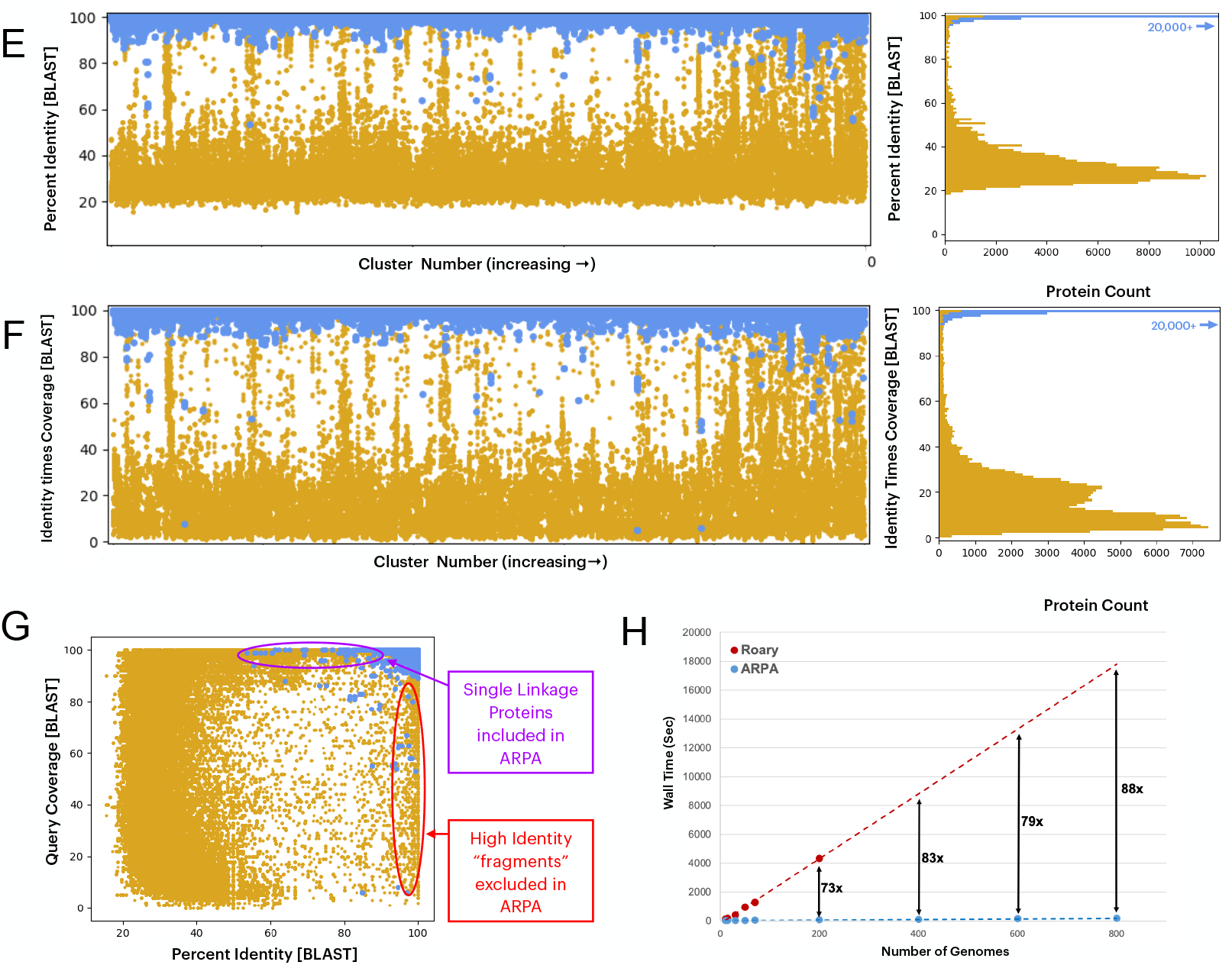
Methodology, Validation and Benchmarking for ARPA clustering. A. Mechanism for vRC clustering analysis. **B.** Number of amino acids in vRCs necessary to identify unique proteins within a *S. aureus* genome **C.** Methodology for single linkage clustering employed in ARPA. ARPA generates two thresholds: (1) proteins that cluster with the initial sequence (default threshold of 95% residue count similarity is user modifiable) and (2) proteins that do not cluster with the first sequence but are similar enough to potentially cluster with other sequences in the initial cluster. This secondary clustering strategy, and further threshold-based analysis ensures that the single linkage clusters are consistent independent of starting vRC. **D.** Use of bSLC method within 50 *S. aureus* genomes reveals that secondary clustering is actively employed when studying biological datasets. **E.** Visualization of percentage identity, **F.** query coverage times percentage identity, and **G** query coverage plotted against percentage identity resulting in BLAST-based confirmation analysis during ARPA validation (blue: sequences within cluster, gold: sequences outside cluster). **H.** Wall time for clustering process with ARPA versus Roary on a personal laptop (Apple Mac 2.6 GHz 6-Core Intel Core i7 with 16 GB RAM). Roary runtimes for runs with large numbers of genomes are linearly extrapolated.

Remarkably, in a phylogenetically diverse survey, as few as 5 amino acid counts from each protein contain enough information to uniquely identify all, or nearly all, proteins in each genome (Fig. 1B). Only two metazoan genomes we tested, *A. thaliana* and *D. melanogaster*, did not follow this pattern likely due to high numbers of isoforms in the input files. Nonetheless, this suggested that in most cases, 20-value vectors could be used reliably to uniquely identify each protein in a genome, and hence, is appropriate for pangenome clustering in many species.

When vRCs are subtracted, the difference represents the amino acid identity between sequences, a value that is largely equivalent to the alignment-based identity statistic and can therefore be used for clustering. Because vRCs allow for tractable one-against-all comparison (see Fig. 1A), we developed a bulk single linkage clustering (bSLC) to cluster homologs based on a 95% threshold (Fig 1C & D). While the time complexity of single linkage clustering is O(n^2^) (Manning *et al*. 2008), bSLC is between O(n^2^), and O(n).

Paralog separation is an additional option in ARPA. Especially in prokaryotes, gene neighborhoods offer a proxy method for paralog separation that is more straightforward than phylogenetic approaches (Esch and Merkl, 2020). ARPA uses homolog clustering as starting information, identifies neighborhoods for all homologs, and splits paralogs when their neighborhoods differ by a threshold (the default is 70% neighborhood identity). While time intensive, this strategy avoids missed paralogous copies that could result from loss of paralogs in genomes.

ARPA outputs three files, two of which are visualizations. The main output is a “.csv” (or “.pickle” for large pangenomics) file, which lists proteins, and a pangenome output table, reporting the presence or absence of proteins and listing an “allele deviation scores” for each protein. This score is calculated as the number of residues different from the single most represented protein sequence within the homolog group.

We tested ARPA clusters using a BLAST strategy. Using an arbitrary protein in each cluster, we compared this protein sequence against all sequences within and outside of the designated protein’s cluster for 10 *S. aureus* genomes. This analysis showed that ARPA clusters can separate sequences with high BLAST identity to the query from other sequences (Fig. 1E). However, because BLAST statistics are based on local not global alignment, percent identity may be very high when only a subsection of a protein is identical, as would be seen in truncated protein. Therefore, we considered query coverage as well (Fig. 1F) revealing that ARPA clustering has 88% sensitivity and 99.8% specificity for identifying BLAST hits that would be considered homologs. The major difference is that ARPA clustering identifies sequences with lower identity but high coverage, while missing high identity low coverage proteins (Fig 1G).

To further test ARPA, we compared it to the widely used application Roary (Page *et al*., 2015). ARPA compares favorably to Roary when run on a personal computer. With clustering, the time advantage generally increases with the number of genomes (Fig 1H). However, the time advantage of this version of ARPA begins to decrease at very large genomic scales. For example ARPA is only 78-fold faster than Roary when analyzing 2,500 *S. aureus* genomes.

Comparing cluster membership of Roary and ARPA for 200 *S. aureus* genomes, we observed high correspondence of the two clustering strategies, including a Rand score of 0.9999 and Adjusted Mutual Information score of 0.9739 (additional metrics in Supplementary Table 1). Additional analysis (see methods; combined cluster analysis) suggested that only 1.1% of clusters do not have a one-to-one correspondence between the two techniques.

Notably, in this analysis ARPA generated more singlet clusters (9130) than Roary (1394) (Supp. Fig. 1.) Comparing the length of the ARPA singlet proteins to the median protein length of proteins within its corresponding Roary cluster (Supp. Fig. 1, inset), revealed that 90% of singlets are more than 5% different in length to the median length of sequence in the Roary cluster. This suggests that the primary difference between Roary and ARPA clustering is the length of sequences and is therefore likely due to insertions, deletions, or truncations.

A maximum-likelihood (ML) phylogenetic inference using the presence/absence of genes as determined by both ARPA and ROARY results in very similar phylogenetic relationships in a test case of 50 *S. aureus* genomes (Fig. 2A). Further, we noted that core genome allele deviation scores within each subclade of *S. aureus* were similar (Fig. 2B). Thus, we hypothesized the allele deviation scores from the core genome could offer a new alternative to presence/absence ML phylogenetic techniques in which the ubiquitous core genome provides no phylogenetic information. Thus, we treated allele deviation scores from ARPA as morphological data in an ML-based analysis for 50 *S. aureus* genomes (Fig. 3A). The topology of the resulting tree is highly similar to the phylogeny from a core gene nucleotide alignment (Fig. 3B, 3C).

**Figure 2.**
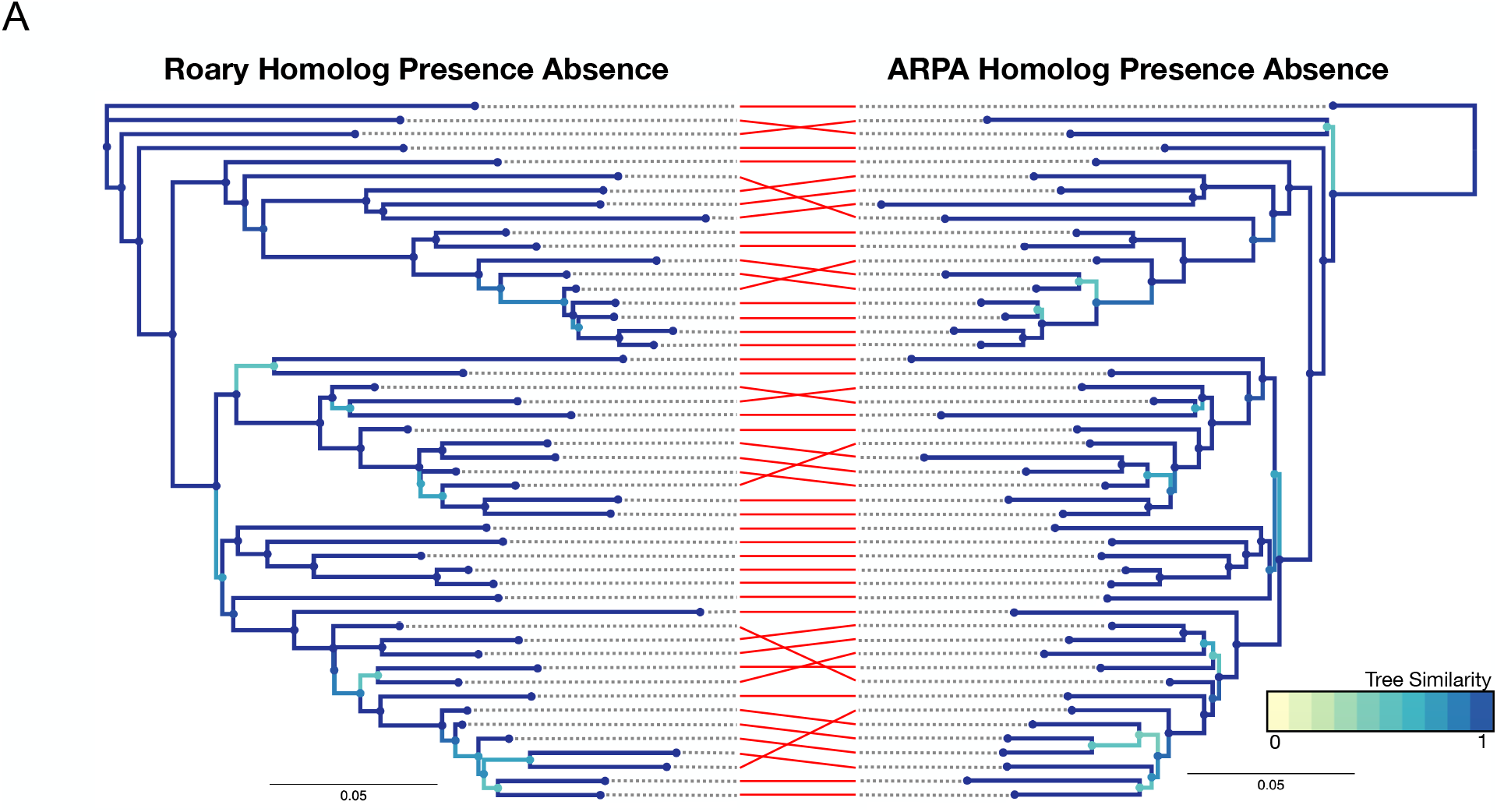
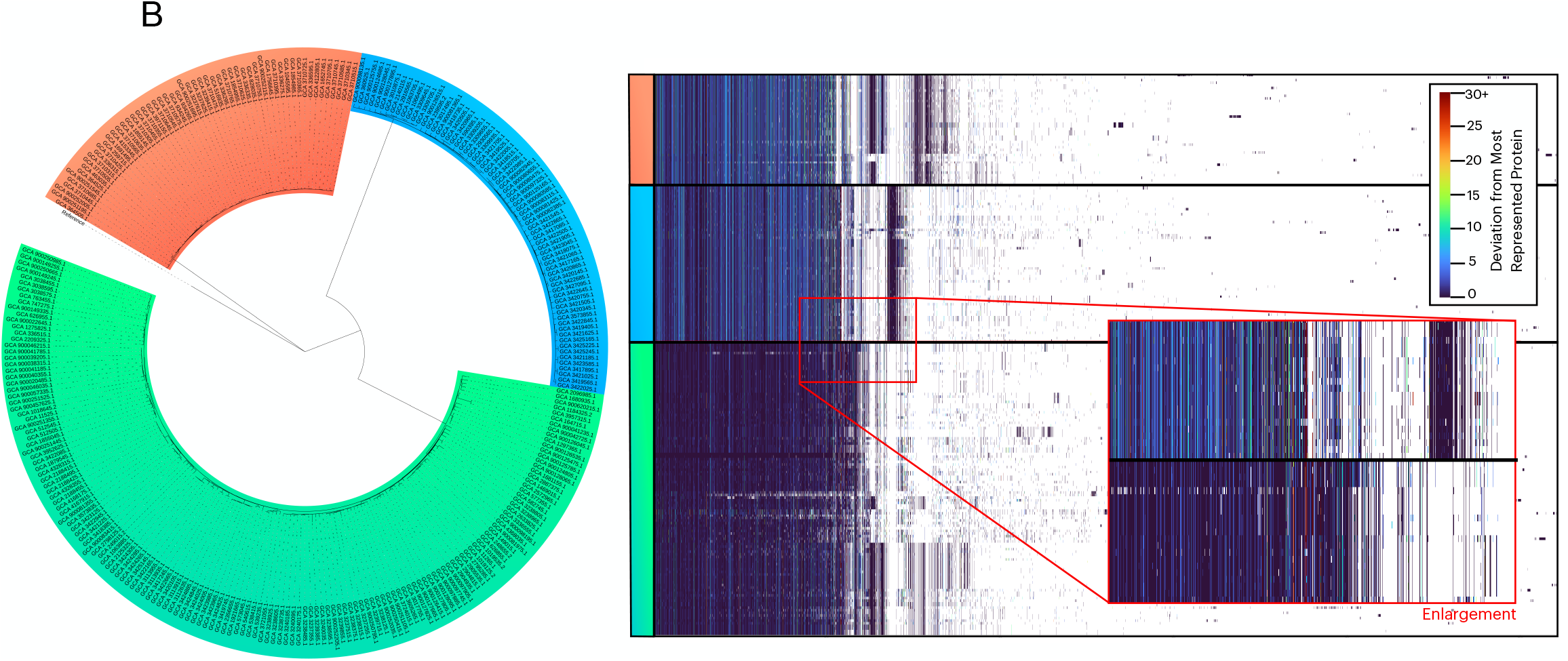
**A.** Maximum likelihood phylogeny and tanglegram of homolog presence absence derived from Roary versus ARPA programs. **B.** Maximum likelihood phylogeny of 218 CC1 genomes, and visualization of corresponding pangenome output from ARPA that demonstrates allelic-level differences in the genomes of species from different subclades.

**Figure 3.**
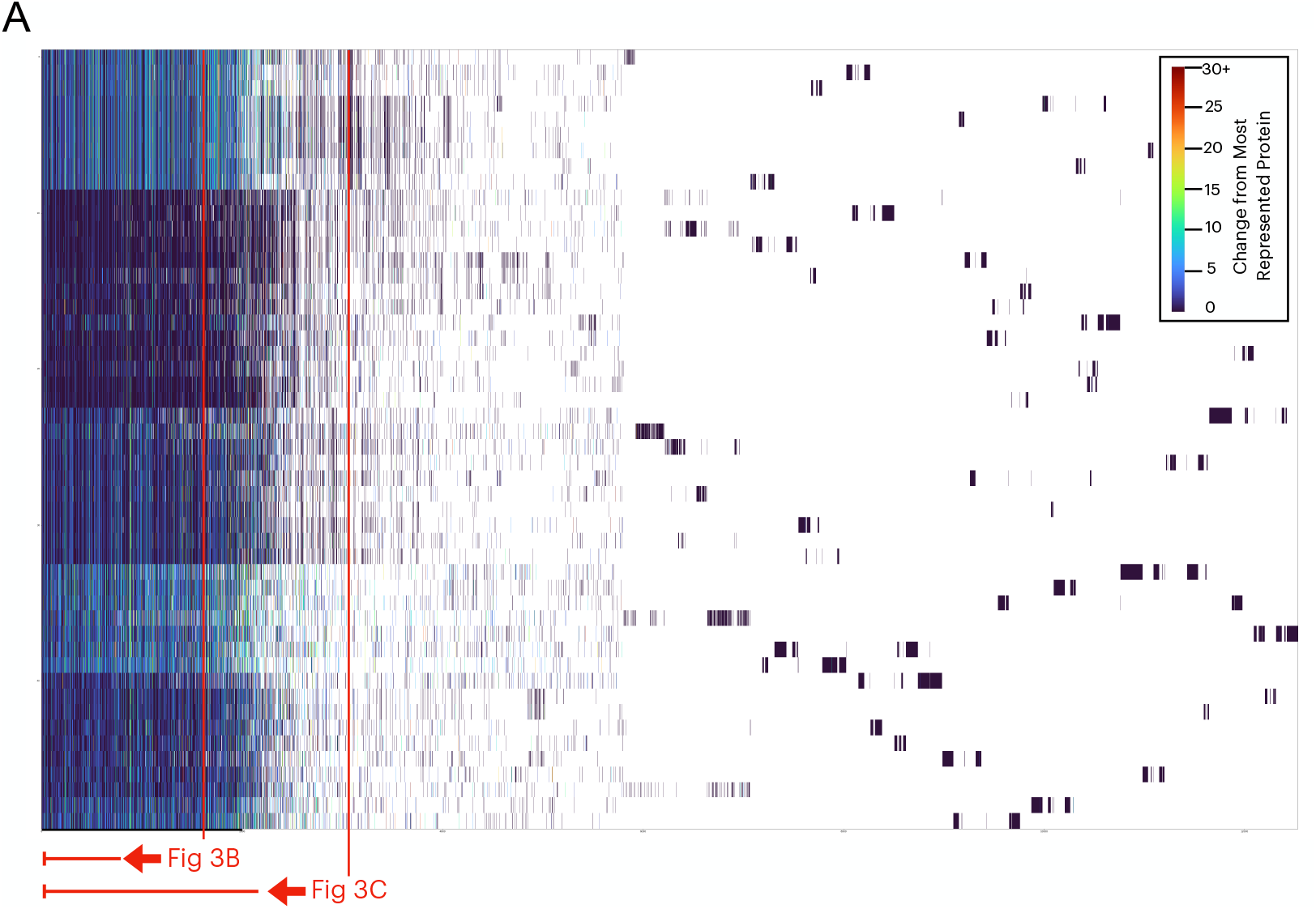
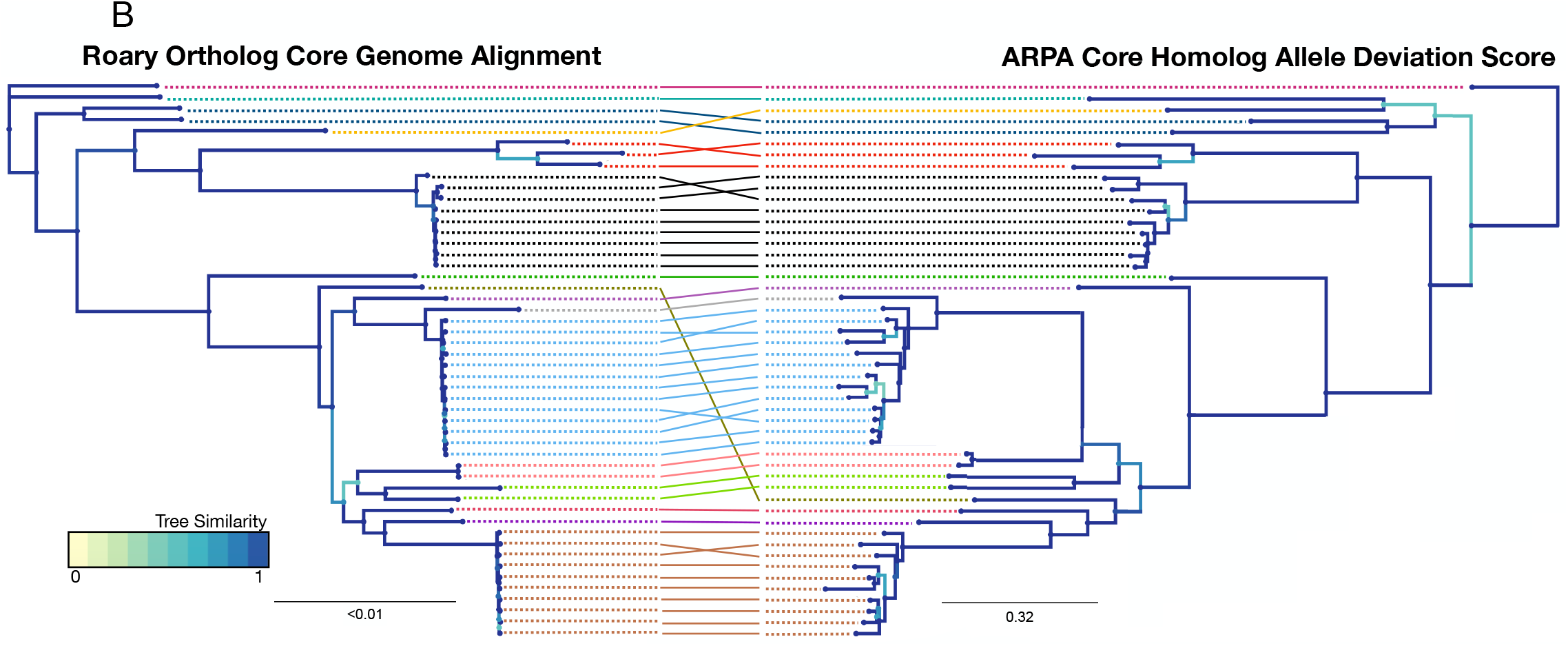
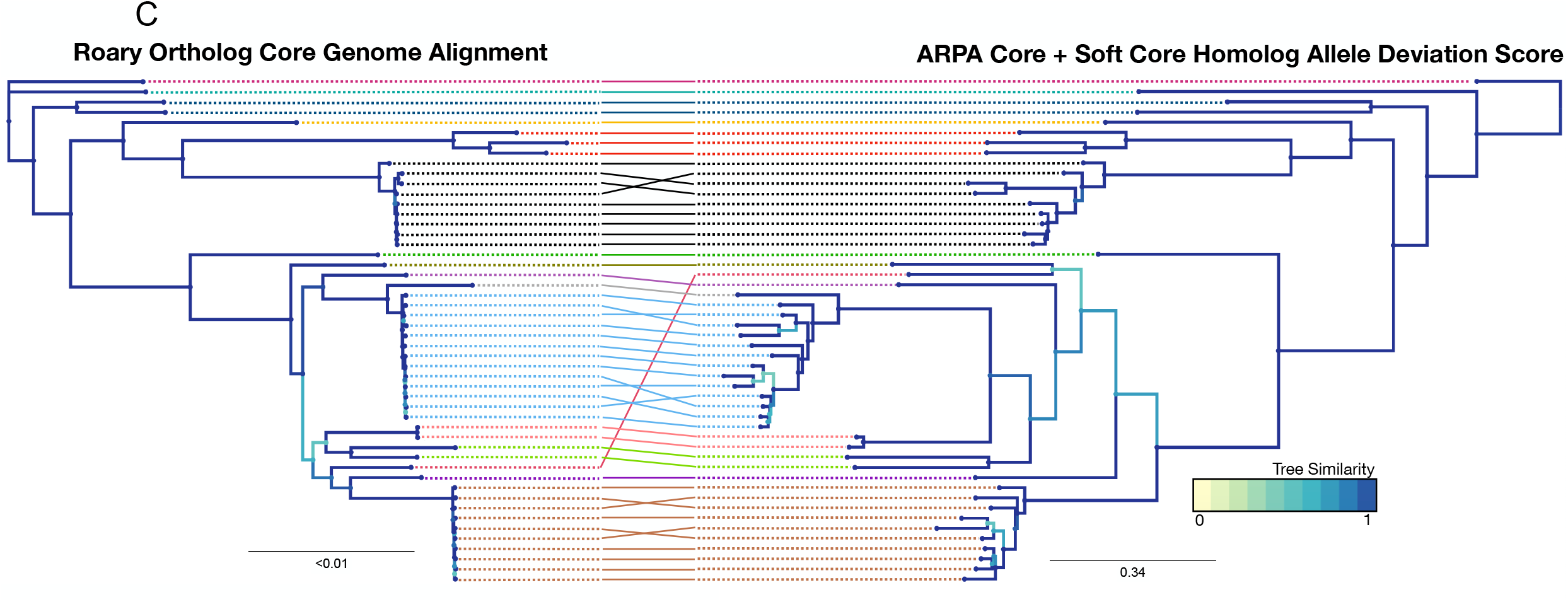
**A.** Visualization of pangenome for 50 *S. aureus* genomes. Phylogeny derived from (**B**) “core” homolog allele deviations and **(C)** “softer core” homolog allele deviations are plotted and compared via tanglegram to a tree derived from core genome alignment. The normalized Jaccard-Robinson-Foulds metric comparing alignment-free trees to the alignment-based tree was 0.24 (Fig. 3B) and 0.20 (Fig. 3C).

The limitations of ARPA are mostly related to the loss of information when sequences are transformed to vRCs. Without alignment some changes will be missed as in the case when two sequences have residues that are swapped in location. As noted before, truncated protein sequences or those with additional sequence may also not cluster using ARPA. It is also formally possible that two distinct, unrelated proteins could have the same combination of residues in different orders. However, our analysis showing that only 5-6 amino residue counts are required to uniquely identify every protein in a genome makes this possibility unlikely in real datasets of closely related organisms. It is also expected that paralog neighborhood splitting will ameliorate this problem.

In conclusion, ARPA is an entirely alignment-free pangenome construction pipeline for pangenomic and phylogenetic analysis that replicates other standard techniques. It has enough computational simplicity to be performed on a regular modern personal computer for thousands of genomes.

## Supporting information

Supplementary Information; NCBI GenBank files used

## Acknowledgments

This work is supported by 1R01AI137526-01 to PJP.

## Methods

Sequences used this manuscript are available from NCBI GenBank, and accession numbers for analyzed sequences within figures are provided in supplementary data.

### ARPA availability and access

ARPA is a publicly available script written in Python that can be accessed through GitHub (https://github.com/Arnavlal/ARPA). Scripts for ARPA verification, as conducted within this manuscript, are available on the GitHub site. The command line interface for this program allows the user to access and edit thresholds dependent upon the specific problem.

### Phylogenetic Tree Building and Analysis

All phylogenetic trees were constructed using IQTree (Nguyen *et al*. 2015). Parameters for each individual tree are listed subsequently in the methods. Phylogenetic trees were visualized through iTOL (Letunic and Bork, 2019). Phylogenetic topological comparisons were visualized with phylo.io (Robinson *et al*., 2016). During comparative analysis, the tree was rotated both computationally and manually, along its nodes to best align in comparative analysis. Tanglegrams were generated manually.

### Internal ARPA analytics

Determination of vRC identification capacity (Fig. 1B) and of the protein count within different aspects of each cluster (Fig. 1D) each were conducted by modifying the ARPA_main.py code to report the numerical data necessary for figure generation. Sequence for vRC residue size protein identification analysis (Fig. 1B) were obtained from NCBI GenBank through accessions: GCA_000001215.4, GCA_000001735.2, GCA_000002415.2, GCA_000005845.2, GCA_000006765.1, GCA_000013425.1, GCA_000146045.2, GCA_000165395.1, GCA_000182965.3, GCA_000195955.2, and GCA_017821535.1.

### Benchmarking with BLAST-based techniques

Clusters from ARPA, after being run with a 90% similarity threshold, were analyzed through BLAST (Camacho *et al*. 2009) within a command line script (“ARPA_Blast_confirm.py”) and analyzed with another script (“ARPA_Blast_Analysis.py”), both available on the GitHub site. Intra-cluster BLAST utilized an e-value threshold of 1e-06. Both within-cluster and out-cluster groups were restricted to only one hit per sequence (“max_hsps”) in order to demonstrate the best alignment of each individual sequence and to prevent the effects of longer sequences from predominating within the analysis. Percentage identity and sequence query coverage were the two metrics reported from this analysis.

### Benchmarking with Roary

Homologous group gene_presence_absence.csv from Roary and its equivalent, modified from the “Pangenome_numerical.csv” for the same dataset were converted manually into a Phylip-like file and analyzed for tree building into IQTree. This analysis used the command “iqtree -s [Filename].phy.” IQTree tested 44 different binary models and selected GTR2+FO+R3 for the Roary dataset and GTR2+FO+R2 for the ARPA dataset as the best-fit models via Bayesian Information Criterion (BIC). 98 candidate trees were analyzed for each tree with 124 and 109 iterations of tree optimization for the Roary and ARPA dataset, respectively. The final model, with optimized state frequencies and site proportions and rates is “GTR2+FO{0.535529,0.464471}+R3{0.76439,0.400663,0.203869,2.30098,0.0317411,7. 07724}” for the Roary Dataset and “GTR2+FO{0.552007,0.447993}+R2{0.882759,0.675289,0.117241,3.44488}” for the ARPA dataset.

Analysis of genomes from the same Clonal Complex (CC1) involved analysis of the pangenome with ARPA. Reference phylogenetic tree for establishing the subclades of CC1 were generated through IQTree. Core genome alignment was performed internally within Roary through MAAFT alignment (Katoh *et al*. 2002) after running Roary, with paralog separation. This alignment was imported into IQTree, which tested 484 DNA models, before selecting SYM+ASC+R2 via BIC. The optimized model after multiple iterations was “SYM{1.03982,3.58997,1.62514,0.3214,3.75392}+FQ+ASC+R2{0.890946,0.857081,0.1 09054,2.16762}”. The phylogeny was visualized within iTOL (Letunic and Bork, 2019). We compared clusters generated in Roary and ARPA using the “Clustered_Proteins” file from Roary and the output file from using the “-check_clusters” option from ARPA. Genomes modified through WhatsGNU_database_customizer.py (Moustafa *et al*. 2020) were used to ensured that genes from “.gff” files in Roary and “.faa” files in ARPA will have identical identifiers. The two resulting text files were imported into a script “ARPA_Roary_clust_confirmation.py,” available on the ARPA GitHub page, where analysis of clustering metrics was done using SciKit package on Python (Pedregosa *et al*., 2011). Scikit run to calculate the adjusted rand score resulted in an error, and so data was exported from Python into R for analysis through the mclust package (Scrucca *et al*., 2016).

Combined cluster analysis occurred through the “ARPA_Roary_clust_confirmation.py” script for 200 *S. aureus* genomes using the “Clustered_Proteins” file from Roary and the output file from using the “-check_clusters” option from ARPA. We scored each protein by its unique membership in each ARPA and Roary cluster, that is if protein X was in ARPA cluster 1 and Roary cluster 1, it was assigned to a cluster A_1_R_1_. If protein Y was in ARPA cluster 1 and Roary cluster 2, it was coded as A_1_R_2_, and assigned to a new cluster. This technique expresses the number of new classifications (A_n_R_m_) when one clustering scheme is combined with another. This method is a metric for cluster correlations; it accounts for the cases where a cluster in one classification scheme is cleanly split into two or more clusters in the other scheme, and when each split cluster maps back to one and only one combined cluster. Therefore, a perfect score, where the two schemes agree completely, will yield a total number of clusters that is equal to the larger number of clusters of the two schemes, in this case a score of 18068 ARPA clusters (Roary generated 6759 clusters). If the number of A_n_R_m_ clusters is higher than this value, then there are some clusters for which there is not a one-to-one correspondence in the other. After applying this to our data, we observed 18271 total groups of cluster classifications, implying that there are 203 (1.1%) instances of clusters that do not have a one-to-one correspondence. Code for supplementary figure 1 analysis and generation are also provided within “ARPA_Roary_clust_confirmation.py” and “ARPA_Roary_singlets.py,” the latter of which requires both the path to clustered proteins files from Roary and ARPA and the original “.faa” file folder.

### Use of ARPA Allele Scores for Phylogenetics

As a reference phylogeny, core genome alignment was performed internally within Roary through MAAFT alignment (Katoh *et al*. 2002) after running Roary and specifying that paralogs should be separated. 484 models were analyzed, with GTR+F+I+I+R4 as the final ideal model for analysis. After 104 iterations of optimization, the final parameters for this tree were “GTR{1.27375,4.55137,1.55387,0.680044,7.16621}+F{0.352358,0.147864,0.192701,0. 307078}+I{0.829107}+R4{0.0477887,2.05068,0.0923509,2.05083,0.0265599,18.682,0. 00419372,51.6039}.”

This phylogeny was compared to the “Pangenome_Numerical.csv” output, with allele differences of up to 30 different states (allele difference scores of 0-29) being preserved within the .phy file conversion (allele differences greater than 30 were counted as missing; this was due to a limit of the number of states possible to specify within .phy morphological file formats). 22 morphological models were sampled, with MK+FQ+R4 selected as the most optimal model for both ARPA core (1617 most represented genes) and soft core (3000 most represented genes). The final “core” model was “MK+R4{0.0960441,0.107221,0.261054,0.50763,0.560738,1. 19508,0.0821638,2.27666 }” after 126 optimization iterations and the final “soft core” model was “MK+R4{0.101485,0.107101,0.241009,0.483032,0.553743,1.14193,0.103762,2.31662}” after 143 optimization iterations.

---

**Supplementary Table 1.** Comparison of cluster performance of Roary versus ARPA using multiple established metrics.

| **Metric** | **Range** | **Output** | **Program Used** |
| --- | --- | --- | --- |
| Rand Score | [0,1] | 0.99995 | Pedregosa *et al.*, 2011 |
| Adjusted Rand Score | [-1,1] | 1 | Scrucca *et al.*, 2016 |
| Normalized Mutual Information Score | [0,1] | 0.98425 | Pedregosa *et al.*, 2011 |
| Adjusted Mutual Information Score | [0,1] | 0.97390 | Pedregosa *et al.*, 2011 |
| V-measure Score | [0,1] | 0.98425 | Pedregosa *et al.*, 2011 |
| Homogeneity (V-measure) | [0,1] | 0.99913 | Pedregosa *et al.*, 2011 |
| Completeness (V-measure) | [0,1] | 0.96981 | Pedregosa *et al.*, 2011 |
| Fowlkes-Mallows index | [0,1] | 0.93711 | Pedregosa *et al.*, 2011 |

**Supplementary Figure 1.**
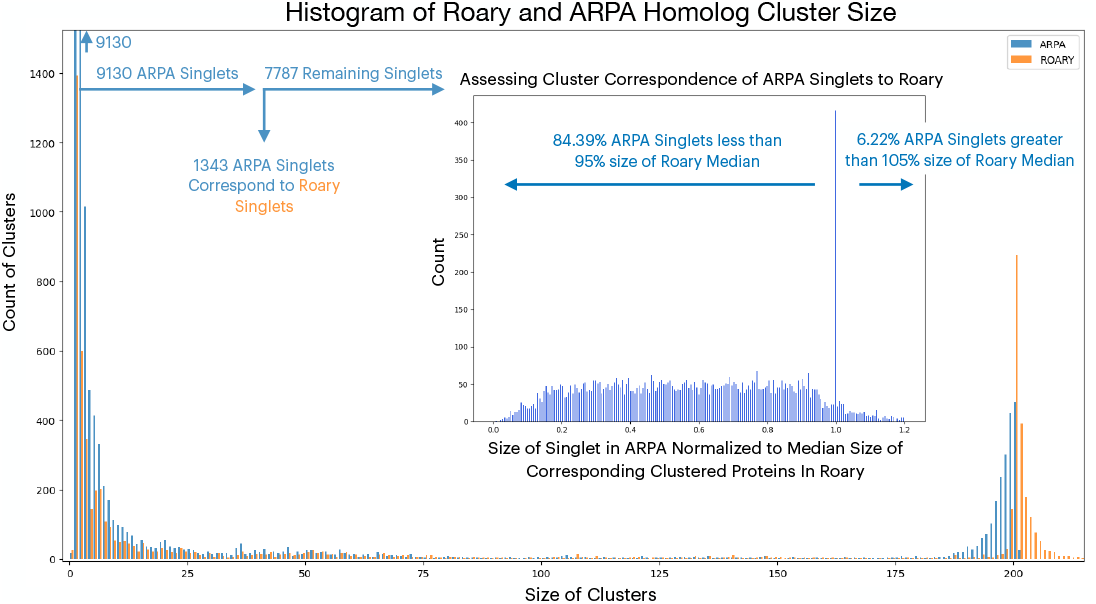
Histogram of Roary versus ARPA cluster size counts for 200 random *S. aureus* genomes. Inset: Determination of the relative size of ARPA singlets versus the median protein size of its corresponding Roary cluster for ARPA singlets that do not correspond to Roary singlets.

